# Medial prefrontal cortex guides goal-directed decision-making about hunger and thirst

**DOI:** 10.1101/620690

**Authors:** Anne-Kathrin Eiselt, Jim Chen, Jon Arnold, Tahnbee Kim, Scott M. Sternson

## Abstract

Decision-making guided by self-evaluation of bodily need states (interoception) is thought to be important for cognitive control over eating or drinking^1^. However, interoception of body states is notoriously unreliable^2–4^ because hunger and thirst have similar motivational characteristics^5^. Consequently, individuals may inaccurately assess their need state and consume food when dehydrated, leading some healthcare professionals to advise overweight patients to drink water before eating^6–8^. Neuroimaging in humans^9–11^ and recordings in rodents^12–15^ indicate medial prefrontal frontal cortex (mPFC) involvement in need state-dependent decisions about hunger and thirst, but mPFC surgical lesions^16,17^ and electrical activity perturbations have little influence on eating or drinking^18,19^. To investigate need-state dependent decision-making as well as the function of mPFC in hunger and thirst, we developed an instrumental foraging task for mice that mimics key elements of human decisions. When homeostatic need state was variable, mice did not show intrinsic knowledge of hunger or thirst state but, instead, rapidly identified their need after consumption of small portions of food and water (outcome evaluation). We observed a food-seeking bias, even in thirsty mice, that required outcome evaluation for mice to correctly seek water. mPFC was required for need state-dependent decisions about hunger and thirst, specifically under variable need state conditions. Food-seeking or water-seeking choices were controlled by multiple decision-making processes, and mPFC was involved in goal-directed decisions about the identity of need states. Thus, we have discovered a role for mPFC in decision-making about hunger and thirst, which is relevant for human behaviors that contribute to obesity.

## Main

Most fluids are consumed during meals, and humans as well as other animals obtain a substantial amount of their water intake from within their food sources^4,20^. Thirst is known to suppress food consumption (e.g., dehydration-induced anorexia^21^) for diets that have very low water content such that the food and food-cues have aversive properties in thirst^22–24^. We were interested in more naturalistic conditions that permit decision-making in hunger and thirst based on physiological need without introducing the aversiveness of dry food in thirst states. To mimic natural food sources that also can be delivered in small quantities during a session, we developed a gelled food formulation with substantial water content (49%) that was minimally consumed in thirst (Fig. 1a, Extended Data Fig. 1a). Preference tests demonstrated that the value of hydrated food during hunger was similar to the value of water during thirst (Extended Data Fig. 1b). To accurately deliver these rewards *via* lick spouts despite the compressible properties of gelled food, we built a novel behavioral apparatus that uses a pneumatic actuator for reward delivery, capacitive lick detection on both reward delivery spouts, and field programmable gate array (FPGA) circuitry for monitoring and control (Fig. 1b, see Methods). We validated our system by confirming that lick-triggered choice of water or gel-food showed strong preferences in thirst or hunger need states, respectively (Extended Data Fig. 1a). Therefore, these conditions show robust contrast in hydrated food and water preference during hunger and thirst without introducing the aversiveness of dry food in thirst states.

**Figure 1:**
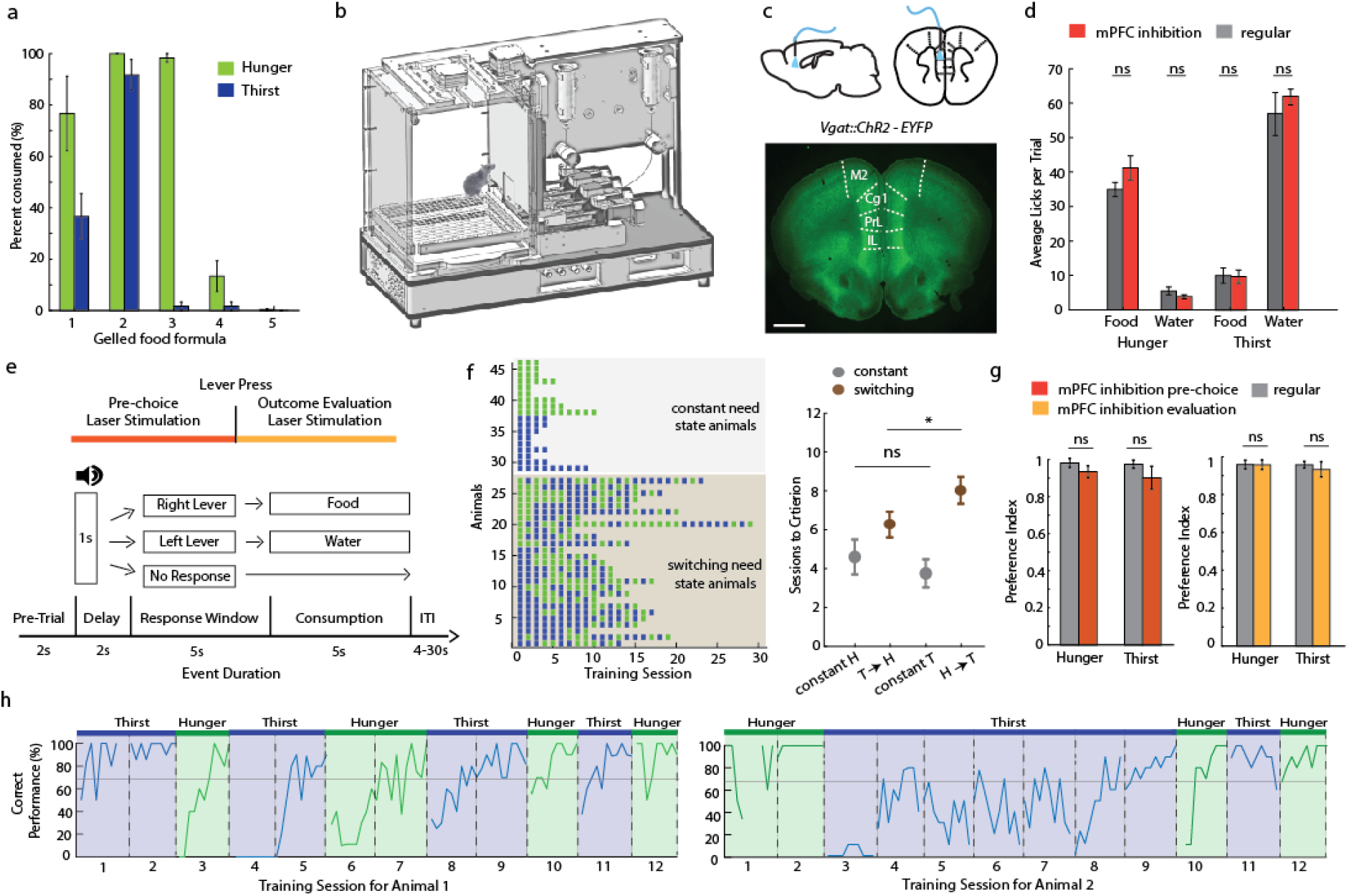
Decision-making about hunger or thirst. **a**, *Ad libitum* consumption of different gelled food formulations in hunger and thirst (n=5). Formulation 3 had the best contrast between the two need states (see Methods). **b**, Diagram of behavioral apparatus. **c**, Sites of optogenetic targeting. Sagittal (upper left) and coronal (upper right) schematic of fiber placement. (Bottom) Coronal section of mPFC from *Vgat::ChR2-EYFP* mouse. M2: secondary motor cortex, Cg1: cingulate cortex area 1, PrL: prelimbic cortex, IL: infralimbic cortex. Scale bar, 1 mm. **d**, Lick-triggered consumption of food and water without (grey) or paired with lick-triggered mPFC silencing (red) in hunger (n=9) and thirst (n=8). **e**, Schematic of behavioral task and timing for optogenetic silencing. **f**, (Left) Individual sessions (green: hunger, blue: thirst) required to reach training criteria for lever presses in constant need state (grey shading) and alternating need state (brown shading). (Right) Mean training sessions in constant need state (grey, n=9 in hunger, n=9 in thirst) and after need state was switched (brown, n=27). **g**, Preference index in hunger and thirst for mPFC silencing during the pre-choice period (left) or reward evaluation period (right) for animals in constant need state (regular n = 13, stimulation during prechoice and evaluation n = 9, for both hunger and thirst separately). **h**, Examples of training performance of need-sate switching animals with low (left) or high (right) food bias in initial training. Testing after each need state switch was separated by 3-4 days. Sessions in the same need state were either consecutive or every other day. Error bars represent SEM. *P<0.05; **P<0.01; ***P<0.001; ns, P>0.05. Statistical analysis in Extended Data Table 1.

We first examined the role of mPFC to guide hunger- and thirst-related consumption behaviors by inactivating local volumes of cortical tissue with millisecond precision by an established method of photostimulating inhibitory interneurons of *Vgat::ChR2-EYFP* transgenic mice^25,26^. mPFC inactivation was targeted primarily to prelimbic cortex (PrL) (Fig. 1c). In closed-loop optogenetic experiments, lick-triggered food or water delivery simultaneously triggered mPFC silencing, which did not affect lick rate or preference for hydrated food or water in either hunger or thirst (Fig. 1d), indicating lack of mPFC involvement in consummatory behaviors.

To investigate decision-making, we developed a two alternative forced choice (2-AFC) instrumental foraging task for obtaining food and water under hunger or thirst (Fig. 1e, see Methods). The task requires similar instrumental actions (lever pressing) to obtain the appropriate outcome (food vs. water) with a similar action for reward consumption (licking) (Fig. 1e). When need state was held constant, mice learned at a comparable rate to correctly press the lever for food or water in hunger or thirst, respectively (Fig. 1f). We calculated a preference index (see Methods) for the outcome appropriate to the need state, which was similarly high in hunger and thirst (Extended Data Fig. 1c). In addition, incorrect choices in thirst that led to food delivery, although uncommon, resulted in food consumption as often as water consumption in hunger (incorrect choice thirst: 27% consumption, incorrect choice hunger: 26% consumption), demonstrating that the incorrect choices have similarly low value in both need states.

The choice between food-seeking and water-seeking requires deliberation about selecting the appropriate action (‘pre-choice’), followed by evaluation of the outcome of the choice (‘reward evaluation’). To dissociate cognitive processes that occur at different phases of the decision, we optogenetically inhibited mPFC during the pre-choice or the reward evaluation periods (Fig. 1e). Silencing mPFC in either trial period did not affect preference index, error rates, average reaction times or average licks per trial in either hunger or thirst (Fig. 1g, Extended Data Fig. 1d-i). Thus, mPFC is not required for making correct instrumental or consummatory choices in mice held in a constant need state.

Because we found little causal influence of mPFC on hunger- or thirst-sensitive behaviors when considered separately, we next investigated the behavior of mice in the 2-AFC task as they were alternating between hunger and thirst states every 3-4 days (see Methods). This is also a more relevant model of the complex physiological dynamics experienced by organisms in resource-sparse environments, in which mice must evaluate their current need state as well as the value of their choices. Mice required significantly more sessions to learn to switch their instrumental response to be appropriate to their need state compared to non-switching (i.e. constant) need state conditions (Fig. 1f) (14.2±1.0 sessions, n=27 vs. 4.1±0.6 sessions, n = 18, mean±SEM, sessions for switching vs. constant need state, respectively; *P*<0.001 Mann-Whitney U test). Moreover, learning the task took significantly longer if mice were initially trained to lever-press in hunger and then switched to thirst compared to animals that were initially trained to lever-press in thirst and then switched to hunger (Fig. 1f,h).

Mice with alternating need states exhibited within-session learning in both hunger and thirst for the correct lever response, such that they had to sample each outcome to correctly guide decision-making (Fig. 2a-c, Extended Data Fig. 2a), even after extensive training (Extended Data Fig. 2b). In contrast, mice in constant hunger or thirst showed high performance throughout each session (Fig. 2c, Extended Data Fig. 2a). For mice switching their need states, cumulative performance was significantly better in hunger compared to thirst (Fig. 2d). This was not due to greater motivation in hunger because lever-press reaction times for water were faster in thirst (Fig. 2e) consistent with prior reports^24,27^, and erroneous responses were significantly slower than correct responses (Fig. 2f). Instead, mice showed a significant food-seeking bias (Fig. 2g) that facilitated correct responding in hunger but was associated with persistence towards food-lever presses at the beginning of thirst sessions (Fig. 2h), which delayed evaluation of water rewards (Extended Data Fig. 2c). In both hunger and thirst, the initial food bias was accompanied by within-session learning of the reward outcomes that were appropriate for the animal’s need state (Extended Data Fig. 2d,e). Thus, under conditions of variable hunger and thirst, mice behaved as if they did not have knowledge of their need state at the beginning of each behavioral session. Instead, mice had a modest bias for food-seeking choices, but they learned to respond correctly for their need by evaluating choice outcome.

**Figure 2:**
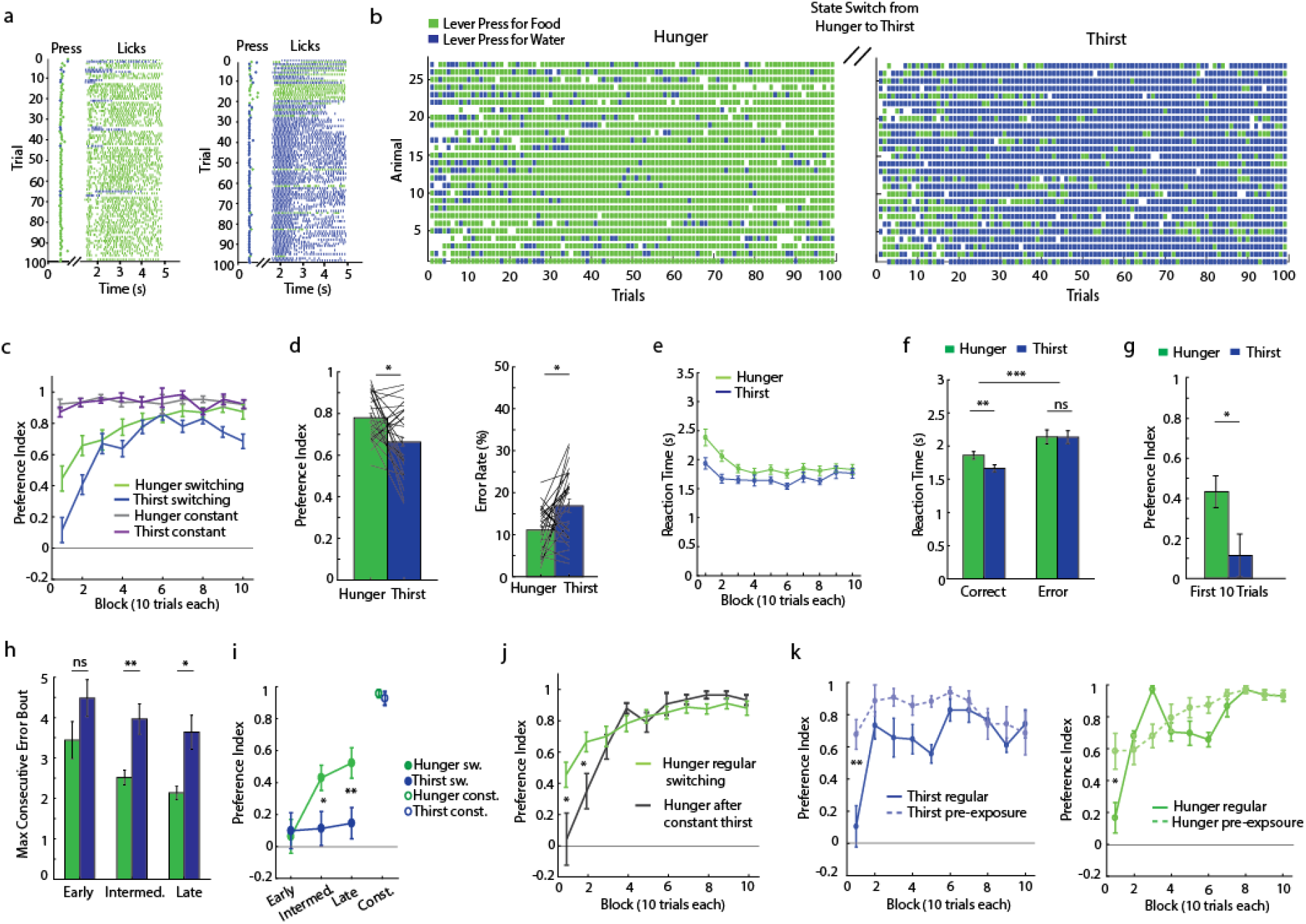
Need-state switching reveals bias towards food-seeking and outcome-driven water-seeking in thirst. **a**, Example of behavioral performance for one animal in hunger (left) and thirst (right), including individual licks per trial (raster). **b**, Decision-making performance of need-state switching for all animals (n=27) in hunger (left) and thirst (right). Choice for food lever: green, or water lever: blue. **c**, Comparison of learning curves for mice in switching (n=27) or constant (n=18) need-state conditions. **d**, Preference index (left) and error rate (right) in hunger (green) and thirst (blue) for need-state switching animals. **e**, Reaction time across session blocks in hunger and thirst (n=27). **f**, Reaction time for correct or incorrect responses (n=27). **g**, Behavioral bias comparison of the preference index during the first 10 trials in hunger and thirst (n=27). **h**, Comparison of the maximum length of consecutive error bouts in hunger (green) and thirst (blue) between sessions during early (n = 27), intermediate (n = 27) or late training (n = 22). **i**, Development of the food bias during early (n = 27), intermediate (n = 27) or late training (n = 22) in comparison with the preference index for constant animals (n= 18). **j**, Comparison of initial food bias during regular switching (green, n=27) and after mice were held in constant thirst and switched back to hunger (black, n=7). **k**, Preference index of regular sessions and sessions with pre-exposure to water and food in thirst (left, n=7) or hunger (right, n=6). Error bars represent SEM. *P<0.05; **P<0.01; ***P<0.001; ns, P>0.05. Statistical analysis in Extended Data Table 1.

Different decision-making processes appeared to underlie the food-seeking bias and the subsequent within-session learning for correct food- or water-seeking responses. The food-seeking choice bias developed after increasing experience with the task across multiple sessions (Fig. 2i, Extended Data Fig. 2a,b). Because short-term consummatory preference tests did not show food-seeking bias (Fig. 1a, Extended Data Fig. 1a), we suspected that this was due to long-term reinforcement by food, possibly reflecting the post-ingestive reinforcing properties of nutrients regardless of need state^28^. Consistent with this, we could eliminate the food-seeking bias, even in highly experienced mice, by maintaining animals in thirst for several sessions followed by a switch to hunger (Fig. 2j). This indicates that the bias towards food-seeking emerges as a dominant choice^29^ in mice frequently switching between hunger and thirst but that this can be controlled by altering behavioral experience to emphasize non-food-seeking actions.

Mice also exhibited prominent within-session learning to achieve correct responding for their need in both hunger and thirst, even after extensive experience with the task (Fig 2c, Extended Data Fig. 2a,b). Fitting the lever press choices to a Weibull distribution^30^ (Extended Data Fig. 2d) showed a significant difference in the offset (Extended Data Fig. 2e), reflective of the initial food-seeking bias, but the learning rate parameters for correct responses across sessions were not significantly different in hunger or thirst (Extended Data Fig. 2e). The similarity of the learning rate parameters indicated an analogous process guiding the improvement of decision-making in hunger and thirst throughout the session. We checked whether mice knew which levers gave food or water outcomes by reversal of lever contingencies, and this led to responding on the lever previously associated with the need state (Extended Data Fig. 2f), indicating that mice knew which levers delivered specific outcomes. Therefore, we investigated the possibility that this learning process reflects uncertainty about the identity of the current need state. Consistent with this, we found that when mice were permitted limited consumption of water and hydrated food in their home cage immediately before transfer to the behavioral apparatus, they significantly improved performance during the initial decision-making trial blocks (Fig. 2k). Thus, once mice determine their need by consuming water and food, they direct their choice to the lever associated with the outcome that reestablishes homeostasis. These experiments suggest that in varying need state conditions, seeking hydrated food becomes a dominant behavioral strategy, but both water-seeking and food-seeking utilize a more cognitively demanding outcome-evaluation process that requires within-session learning.

Next, we investigated the role of the mPFC in decision-making under conditions of variable need. Silencing mPFC in the pre-choice period of the trial greatly reduced performance in thirst (Extended Data Fig. 3a-c). Strikingly, most mice (13/17) incorrectly pressed for food and consumed food rewards in thirst (Fig. 3a, Extended Data Fig. 3a, Movie 1, see Methods), which typically, but not always, returned to correct responding in trial blocks lacking mPFC inactivation. Inactivating the same cortical area in the same mice during hunger did not affect decision-making (Fig. 3b,d, Extended Data Fig. 3a-c). Reaction times for correct presses were significantly slower for stimulation trials in both hunger and thirst (Extended Data Fig. 4a), but lick rates following the choice were unaffected (Fig. 3d). Thus, mPFC guides correct responding for thirsty but not hungry mice under conditions of variable need states.

**Figure 3.**
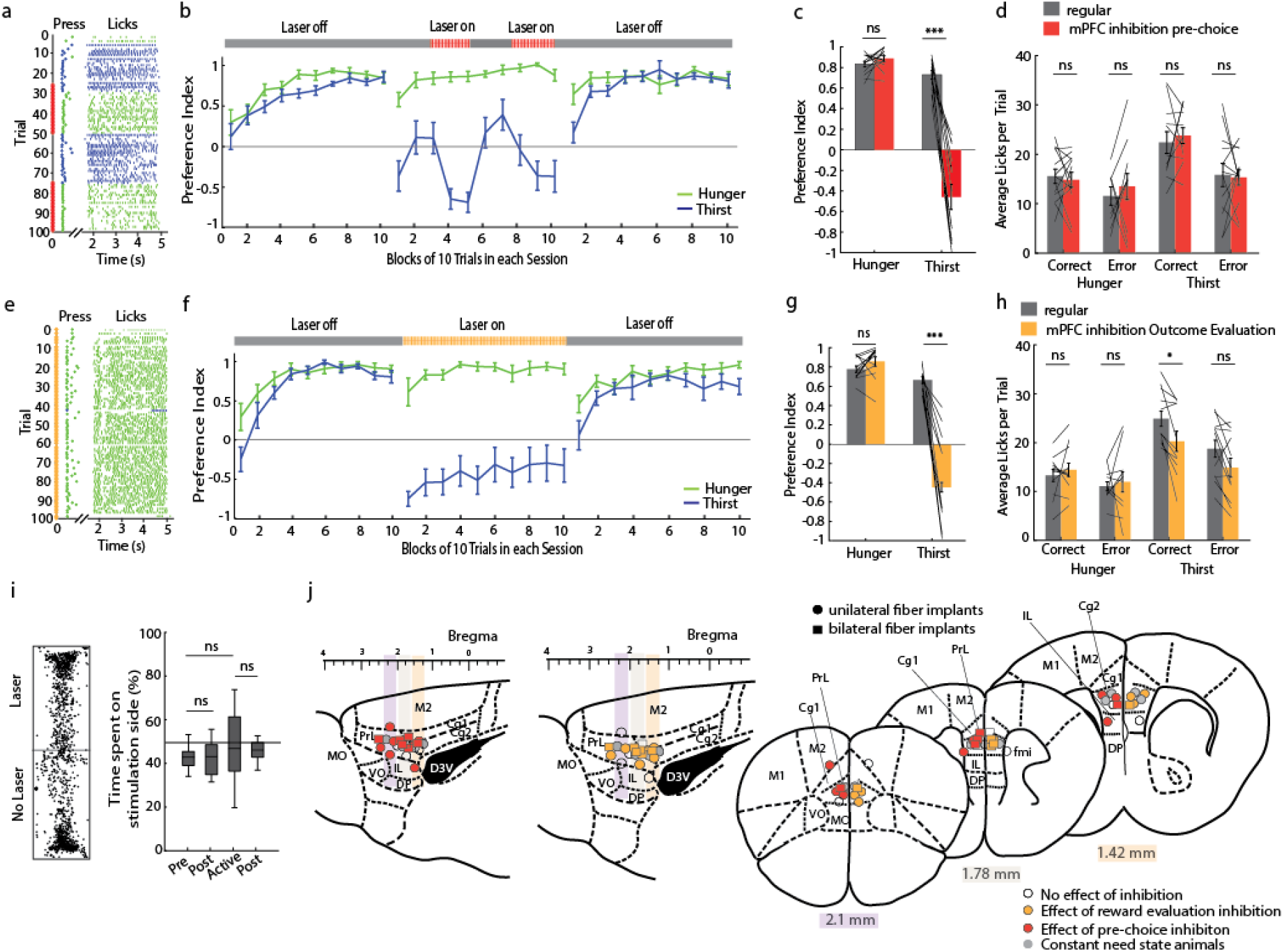
Optogenetic inhibition of mPFC in thirst drives incorrect food-seeking. **a**, Example session performance in thirst with pre-choice mPFC inhibition. **b**, Mean performance in hunger and thirst during three separate sessions: before, during and after pre-choice mPFC inhibition for all mPFC-sensitive animals (n = 13, see Methods and Extended Data Fig. 3 for all subjects). **c**, Preference index comparison between pre-choice mPFC inhibition (red) and regular responses (grey) in hunger and thirst in the same trials (trial 25-50 and 75-100). **d**, Effect of mPFC inhibition during pre-choice period on licks per trial. **e**, Example session in thirst with mPFC silencing during reward evaluation period. **f**, Mean performance in hunger and thirst during three separate sessions: before, during and after reward evaluation mPFC inhibition for all responding animals (n = 12, Extended Data Fig. 3 for all subjects). **g**, Preference index comparison between reward evaluation mPFC inhibition (orange) and regular responses (grey) in hunger and thirst during the whole session (all 100 trials). **h**, Effect of mPFC inhibition during reward evaluation period on licks per trial. **i**, Conditioned place preference test (n=12). **j**, Anatomical location of optical fiber placement. Colored sites are locations that elicited a deviation from the mean of regular performance by at least 2 standard deviations with inhibition during the pre-choice period (red) and reward evaluation period (orange). Fiber placement in constant need state animals is depicted in grey. Cg1: cingulate cortex area 1, Cg2: cingulate cortex area 2, DP: dorsal peduncular cortex, D3V: dorsal 3rd ventricle, fmi: forceps minor of the corpus callosum, IL: infralimbic cortex, M1: primary motor cortex, M2: secondary motor cortex, MO: medial orbital cortex, PrL: prelimbic cortex, VO: ventral orbital cortex, Error bars represent SEM. *P<0.05; **P<0.01; ***P<0.001; ns, P>0.05. Statistical analysis in Extended Data Table 1.

Because mice that are repeatedly switching between hunger and thirst show an initial period of learning at the beginning of each session, we also investigated the effect of inactivating mPFC only during the reward evaluation period. Neither unilateral nor bilateral silencing of mPFC after the decision affected choices in hungry mice, but thirsty mice (12/17) were profoundly affected, with many mice completely reversing their choice to food-seeking throughout the entire session (Fig. 3e-h, Extended Data Fig. 3d-f, Extended Data Fig. 4b). These are amongst the most robust effects of mPFC perturbations on need-based behaviors ever reported, and it is notable that the perturbation was after the choice. This was not due to the valence of mPFC photoinactivation, which did not influence place preference (Fig. 3i). Consumption behavior during mPFC inhibition was unaffected in hunger, but slightly decreased in thirst (Fig. 3h), which is likely related to inappropriate consumption of food in thirst. Subsequent sessions in thirst without modulation of the mPFC showed normal water-seeking performance (Fig. 3b,f and Extended Data Fig. 3a,d). Sensitivity to mPFC silencing in the reward phase with thirsty mice was tightly circumscribed within the PrL and adjacent rostral anterior cingulate cortex (rACC), with most non-responders on the periphery or outside of this region (Fig. 3j, Extended Data Fig. 5). Thus, mPFC is critical for the evaluation of behavioral choices in mice under conditions of homeostatic variability.

**Figure 4.**
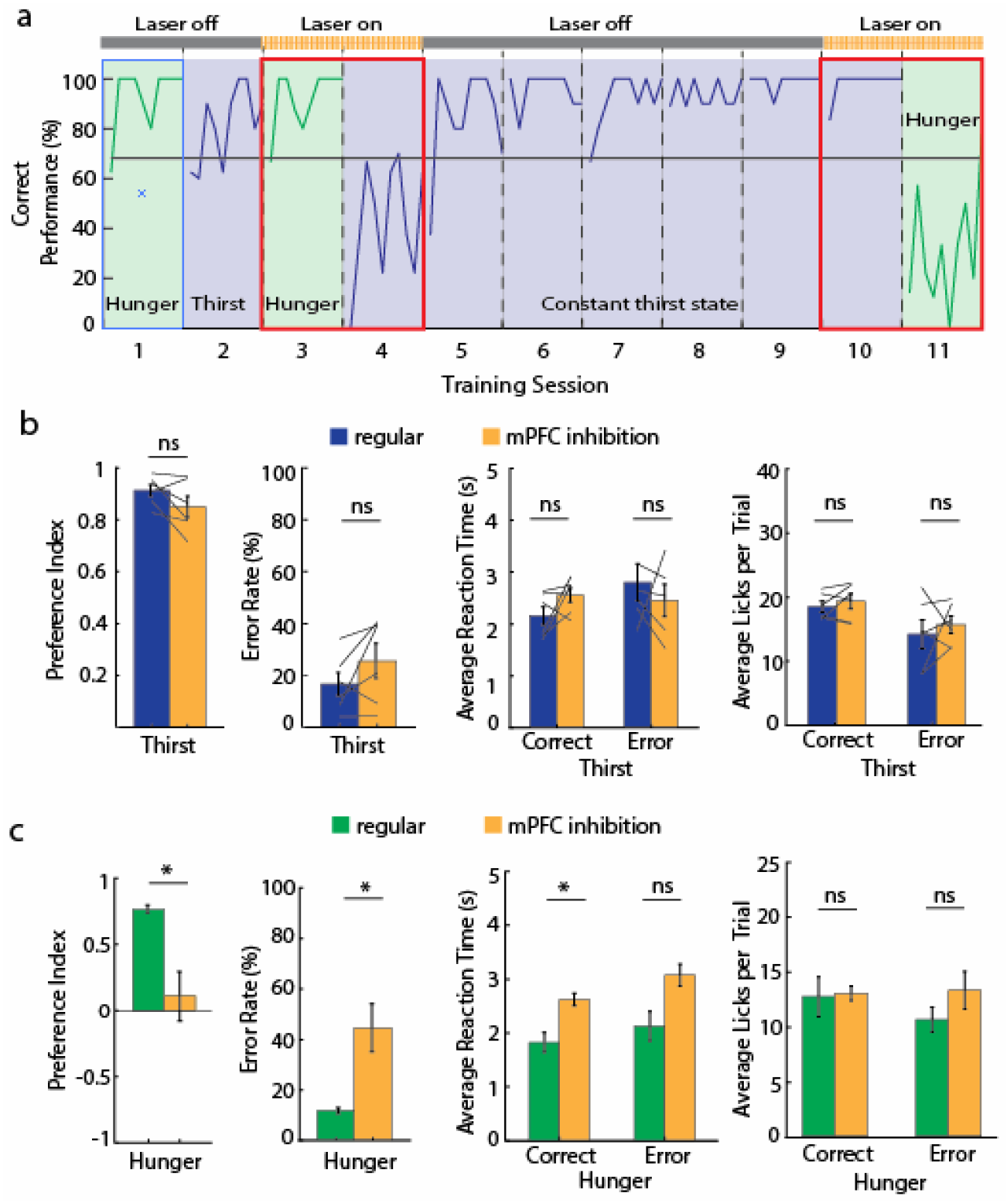
mPFC controls evaluative decision-making in both hunger and thirst. **a**, Example lever press performance for one animal during need-state switching with mPFC silencing during reward evaluation in hunger and thirst before and after constant thirst state. **b**, Preference index, error rate, reaction time and licks per trial in thirst during regular need-state switching (blue) and with mPFC silencing during reward evaluation after holding thirst state constant for several sessions (orange) (n = 6). **c**, Preference index, error rates, reaction time and licks per trial in hunger after being in constant thirst for several sessions without (green) and with mPFC silencing during reward evaluation (orange) (n = 7). Error bars represent SEM. *P<0.05; **P<0.01; ***P<0.001; ns, P>0.05. Statistical analysis in Extended Data Table 1.

Most mice that were affected by mPFC silencing showed sensitivity during both the choice and the reward phase of the behavior (Extended Data Fig. 5a), however this was not the case for all animals. This indicates that different but overlapping neuronal networks are engaged by different phases of the decisionmaking process in mPFC. Moreover, prefrontal cortex silencing had no effect on choice behavior in hunger. Therefore, decision-making in thirst, but not hunger, relies on the mPFC and inhibiting this area led to a default food-seeking behavior that was also evident in the biases we observed when training animals with alternating need states (Fig. 1f).

The reliance on mPFC during thirst but not hunger suggested either a specialized role of this brain region in thirst or it could reflect selective involvement of mPFC in the outcome evaluation decision-making strategy that was especially prominent during thirst. Furthermore, we suspected that the dominant behavioral strategy^31^, usually food-seeking, was independent of mPFC and was implemented as the default response when mPFC was inhibited. We tested this hypothesis by transitioning mice trained under need state variability to a constant thirst state for several sessions (Fig. 4a), which we have shown eliminates food-seeking as a dominant behavioral response (Fig. 2j). Mice that were previously sensitive to mPFC silencing and switched their behavior to food-seeking in thirst, now showed no reduction of water-seeking behavior during mPFC inactivation (Fig. 4a,b). Under these conditions, mice adopted a new dominant behavioral response of water-seeking, which was now independent of mPFC function.

We predicted that if mPFC function is related to learning need state identity under variable homeostatic conditions, then food-seeking in hunger would now be mPFC dependent because it was no longer the dominant behavioral response. Indeed, after consecutive thirst sessions, when mice were switched to hunger, their decision-making became sensitive to mPFC inactivation and exhibited pronounced water-seeking in hunger (Fig. 4c).

Taken together, these experiments identify an important role for the mPFC in decision-making under variable need states by evaluating decision outcome. In contrast, mPFC did not affect dominant response strategies associated with constant need or a habitually favored outcome. Previous reports indicated that rodents store different outcome values in long-term memory for hunger and thirst, which can guide correct responding after a single need state switch^31^. Our frequent switching between hunger and thirst leads to lower initial decision-making performance and indicates limitations on cognitive awareness of those need states in mice. Nevertheless, mice quickly achieved high performance within a session if they were pre-exposed to the rewards, demonstrating that frequent need state switching leads to reliance on outcome evaluation to guide decision-making. Our results are consistent with a role of the mPFC in incentive learning^32^ (Fig. 3f), as well as its involvement in guiding action selection (Fig. 3b)^29^.

We also found that the use of a food source with ethologically relevant water content led to a progressively learned choice-bias towards food-seeking in mice, which may contribute to food-seeking biases in humans that can lead to obesity. The clinical suggestion to drink water before meals effectively promotes a water-seeking habit^7^ and, based on our results modeling this treatment (Fig. 2j), it may reduce food-biased choices in thirst and aid weight-loss management^8^. In principle, other non-food habits may also reduce the bias towards food-seeking behavior. This has been shown to be more effective than simply promoting a reduction in food-seeking behavior without replacing it with another behavioral goal^33^. Although additional work is needed to dissect the underlying cellular and molecular mechanisms in the mPFC, our findings are the first to reveal a high order neurobiological substrate in the prefrontal cortex that is required to properly guide decision-making between water- or food-seeking in hunger and thirst.

## Acknowledgements

This research was funded by the Howard Hughes Medical Institute. We thank S. Michaels for histology; M. Rose, M. McManus, R. Gattoni, S. Erwin, C. Morrow, A. Zeladonis, C. Lopez for mouse breeding and procedures; A. Hantman, J. Dudman, U. Heberlein, A. Hermundstad, R. Egnor for comments on the manuscript.

## Author Contributions

A-K.E and S.M.S initiated the project, designed experiments, analyzed the data and prepared the manuscript. A-K.E, J.C. and J.A. designed, engineered and tested the behavioral apparatus. A-K.E. performed experiments and collected data, T.K. performed experiments and collected data.

## Methods

### Mice

Adult male and female (over two month old) *Vgat::ChR2-EYFP* transgenic mice (Jackson laboratory, n = 37) were included in all behavioral and photostimulation experiments. Male and female (over two month old) C57BL/6J mice (Jackson Laboratory, n = 10) were used for behavioral experiments. Mice were individually housed in a temperature- and humidity-controlled room and maintained on a 12-h light/dark cycle. All animals were handled according to US National Institutes of Health guidelines for animal research and experimental protocols were approved by the Institutional Animal Care and Use Committee at Janelia Research Campus.

### Gelled Food

We tested several different gel food mixtures with varying degrees of water content (ranging from 30-60% water content) and different thickening agents. All mixtures were made of water and standard dry powdered food mix from TestDiet (PMI Micro-stabilized rodent liquid diet LD101, www.testdiet.com), a nutritionally-balanced, easy to prepare powder that contains 17.7% protein, 16.9% fat, and 65.4% carbohydrates, and minerals. The five gel food mixes we tested in the free consumption test in hungry and thirsty mice (Fig. 1a) contained water (49%), food (50%), and up to 1% of the following thickening agents: Formula 1, gelatin; Formula 2, Thicken Up (a baby food thickener with xanthan gum from Nestle Health Science); Formula 3, Carrageenan; Formula 4, Xanthan Gum; Formula 5, Cornstarch. We used Formula 3, because mice showed low consumption of this mixture in thirst but consumed all of it when hungry. To create this gelled food mix, dry powdered food (10 g) was mixed with distilled water (10 ml) and the food thickening agent, Kappa Carrageenan (0.15 g). All three ingredients were thoroughly mixed and centrifuged for 8 minutes at 1200 rpm at 4°C to eliminate air content, reduce compressibility and ensure precise and consistent partitioning of the gel food using the solenoid valve and pneumatic system.

### Food and Water restriction

Mice were kept on food or water restriction with daily health monitoring and body weight assessments. Restriction was eased if mice fell below 70% of their initial *ad libitum* fed body weight or failed qualitative health assessment. For water restriction, mice received approximately 1 mL water daily, with ad libitum access to rodent chow (PicoLab Rodent Diet 20 5053, www.labdiet.com, water content 10%) in the home cage. For food restriction, mice received 2-3 g of food pellets in their home cage, with ad libitum access to water. In constant need state conditions, mice were held in either hunger or thirst, unless otherwise noted, and experiments took place every 2-4 days. For need state switching conditions, animals were switched from food restriction to water restriction immediately following the experiment and, after behavioral testing in the thirst state, mice were subsequently switched from water restriction to food restriction. For these cycles of hunger/thirst switching, we allowed 3-4 days between switching need-state experimental sessions to ensure that animals were sufficiently restricted in each state.

### Behavioral apparatus

We engineered a system with four motorized slides that could be extended into or retracted out of the behavioral cage. Two of the slides held levers with limit switches that could be pressed by the animal, while the other two slides held tubes that could dispense food or water reward. Lick detection on the tubes was performed by capacitive sensing. Food and water were dispensed using solenoid pinch valves (NResearch Corporation). Water was gravity fed from a syringe reservoir while the gelled food mixture was dispensed from a syringe whose plunger was driven by air pressure. Food and water dispense volume requirements were 6 ± 1.5 μl. Slide motor control, sensor data collection, pulse sequence for laser, video display and dispense control were all controlled with a custom FPGA control board. The FPGA control board also logged imaging and sensor data. Behavioral video images were logged at a frame rate of 30 Hz. Each image frame is processed on the FPGA and the frame embedded with the collected sensor data. The final image is then sent to a connected control PC over a 5Gbps Cameralink port. The control PC runs software written in C/C++ with a user configurable state machine for cage control. The experiment parameters are set using the control software GUI on the PC and sent via UART to the FPGA. The PC then extracts the sensor data from the images and stores the data separately on the hard disk.

### Free consumption choice task

For the free consumption choice task, as well as for all behavioral experiments reported here, we chose a reward size of 6 μl water and 6 μl gelled food so animals would reliably perform over 100 trials per session and not be satiated within that time frame. To test the rewarding properties of the water and food reward, we used a lick-triggered consumption task, where mice had access to the water and food spouts for 15 s each trial and every 10^th^ lick at either spout resulted in the delivery of a food or water reward. After 15 s both spouts were retracted, followed by an inter-trial interval between 4-30s. Each session consisted of 50 trials. The average amount of reward delivered in that 15 s time period was similar in hunger and thirst, suggesting that the intensity of the need state and the value of the rewards were comparable.

### Behavioral task

Mice were handled and acclimatized to the behavioral cage for at least one day and were then either trained in the 2-alternative forced choice task (2-AFC) or tested in the free consumption choice task before being trained in the 2-AFC task. In each trial of the 2-AFC task, the presentation of a tone (1 s, 12 kHz) indicated trial onset. After a delay (1 s), both levers were extended and available (5 s response window). Pressing the one lever delivered the food reward (6 μl), whereas pressing the other lever delivered water (6 μl). The lever-reward contingencies were randomized across animals, meaning that for half of the animals the right lever was associated with food and the left with water, whereas for the other half the right lever was associated with water and the left with food. There was no difference in performance or reaction time between the two different lever-reward contingencies. After a lever press was detected, both levers retracted, and the reward spout associated with the pressed lever was extended. Animals had access to the reward spout (5 s consumption window) before it was retracted followed by a variable inter-trial interval (4 – 30 s). If no press occurred during the response window, levers were retracted, the inter-trial interval began, and the behavioral response was counted as a miss. Animals completed 100 trials in each session. A session was considered successful, if the animal pressed at least 68% for water in thirst or food in hunger.

### Behavioral training

During the initial training period the response and consumption window was 20 s, which were gradually reduced to 5 s throughout the following training sessions. If mice pressed more than 68 presses for the correct need the session was considered successful. Mice in switching need state conditions were switched to the other need state, whereas constant need state mice stayed in one need state and were trained or tested every 2-4 days. All constant need state mice had at least 4 successful sessions before mPFC optogenetic silencing experiments. Mice undergoing need-state switching were tested in their new need state 3-4 days after the switch and had to learn to press the opposite lever to receive the reward appropriate to that need state. If mice successfully pressed for the appropriate outcome, the need state was switched again. This continued until mice responded with more than 68 presses for the correct outcome in 3 consecutive switching sessions. If the performance did not reach that criterion, an additional session in the same need state took place until that criterion was reached and the animal was again tested with switching states until 3 consecutive sessions reached that threshold. The range of training time for animals alternating their need states was 3-9 weeks. After completion of the initial training, all experimental sessions took place, typically a duration of 16-20 additional weeks. If the animals’ performance was affected by optogenetic mPFC silencing or any other manipulation and thus did not reach 68% correct, a regular session in that state was performed the next day before the need switch to ensure retention of the task.

### Pre-Exposure task

For the pre-exposure task, we provided mice with a small amount of both water and food into the homecage immediately before the onset of the lever pressing session. The amount provided equaled what animals could earn during the first 10 trials pressing the food or water lever: 60 ul water and food. All animals consumed most food and water provided in the home cage, irrespective of their need state (hunger vs. thirst).

### Reversal task

To test the hypothesis that the initial within-session learning was due to uncertainty about which lever gives the food or the water reward, we reversed the lever contingencies. Mice continued switching between need states, but both levers gave the opposite outcome that was originally associated with that lever (i.e., the former food lever now delivered the water spout and the former water lever delivered the food spout). If mice used a behavioral strategy in which they determine in each session which lever gives the greatest reward for their current need, then reversal of the lever contingencies should not affect performance. Instead, we found that the initial performance was opposite to the reversed contingencies (Extended Data Fig. 2f). This indicates that in both hunger and thirst, mice were seeking the outcome appropriate for their need, which led to the incorrect responses due to the reversed contingencies.

### Fiber implantation and optical stimulation

Fiber implantation surgery was performed under anesthesia (1.5% isoflurane). The skull was exposed and customized fiber optic probes (200 μm diameter core, multimode, NA 0.48, ThorLabs) were implanted above the mPFC either unilateral (coordinates from bregma: −1.8mm to −2.0mm AP; −0.2mm to −0.5mm ML; −1.3mm to −1.6mm; 3–5° angle) or bilateral (coordinates from bregma: −1.8mm to −2.0mm AP; ±0.5mm to ±0.8mm; DV −1.5mm to −1.9mm; 5−12° angle). Animals had at least 10 days to recover from surgery before food and water restriction and training began.

Experiments involving optogenetic activation of cortical interneurons in *Vgat::ChR2-EYFP* mice expressing the light sensitive cation-conducting opsin, channelrhodopsin-2 in GABAergic interneurons were performed using λ = 473 nm blue light at 6-8 mW laser power at the tip of the fiber with 10 ms pulses of light at 20 Hz frequency. Light was delivered in two different time periods: (1) the pre-choice period starting at the onset of the cue until a lever was pressed, or (2) the outcome evaluation period, which started after the lever was pressed until the reward spouts were retracted. For the pre-choice period, stimulation took place from trial 25-50 and trial 75-100, while trial 1-24 and 51-74 were laser off conditions. For the outcome evaluation period, all trials (1-100) were laser-on trials to prevent regular, undisturbed outcome evaluation during the session and, thus, prevent within session learning about the value of the outcome.

### Histology, immunohistochemistry and microscopy

Animals were anaesthetized with isoflurane and transcardially perfused with PBS followed by 4% paraformaldehyde in PBS. Brain sections (50 μm) were imaged to determine fiber placement on an upright epi-fluorescent microscope with 10× or 20× objectives.

### Conditioned place preference

Conditioned place preference was performed as previously described^5^. In brief, a sound-isolated, two chamber apparatus with visual and textual distinct sides was used and an overhead video camera recorded the position of the animal. After acclimatization, hungry or thirsty animals were placed in the apparatus for 30 min and their initial preference was recorded. The less preferred side was then paired with photostimulation for 30 min with 10 ms pulses at 20Hz for 1 s, repeated every 4 s in a passive conditioning task for 5 consecutive days. On the same 5 days but at different times of the day and at least 5 hours apart, animals were tethered and placed on the preferred side for 30 min but without receiving photostimulation to match the time spent on each side of the chamber. After that preference was tested again. We also performed a close-loop place preference in the same animals, in which they had access to both sides of the chamber and photostimulation was applied when the mouse entered the less preferred side (which was previously paired with the passive conditioning stimulation). Photostimulation ceased as soon as the mouse crossed to the other side. The next day free access preference was tested again.

### Data analysis

#### Preference index

The preference index (PI) was calculated subtracting incorrect presses from correct presses and dividing by the number of total presses.

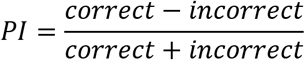

For the learning curve, we calculated the PI for a block of ten trials throughout the session for a total of 10 blocks (100 trials).

#### Transition trial and maximum error bouts

The transition trial was calculated with a sliding window analysis (window size of 10 trials, step size 1 trial), until the animal reached 80% correct responses (8 correct presses out of 10 total presses). To derive the maximum length of error bouts, we calculated the number of consecutive errors and chose the longest bout in each condition.

#### Reaction time analysis

We excluded all missed trials (trials where the animal did not press either lever) from the analysis and calculated the mean reaction time for all presses for the water and food lever in hunger and thirst. Trials in which animals reached outside the cage to press the lever (i.e. before the lever was fully extended into the behavioral cage) were included in the analysis.

#### Lick Analysis

For the free *ad libitum* consumption choice task, both spouts were exposed and animals could lick both spouts. Total number of licks on each spout was recorded and averaged for each animal in hunger and thirst. If no licks occurred during any given trial it was counted as value 0 and included in the analysis. For the instrumental lever pressing task, average licks per trial were calculated for each animal for all trials where the food or water lever were pressed and the food or water spout was exposed for consumption of the reward. Trials where animals did not lick from the spout were assigned with the value 0 and included in the calculation.

#### mPFC silencing responders versus non-responders

For the optogenetic silencing experiments, we report the data for responder mice that showed an effect of laser stimulation greater than 2 standard deviations above or below the mean error rate of 3 independent non-stimulation sessions in both hunger and thirst. The same mice were used in both hunger and thirst conditions and thus differences in sensitivity to mPFC silencing in hunger and thirst are measured across the same animals. We choose to display the effect of mPFC silencing for responders only (Fig. 3, Extended Data Fig. 4, and Extended Data Table 1) to emphasize the distinct sensitivity to mPFC silencing from thirst sessions to hunger sessions. Inclusion of all mice in the grouped analysis also shows significant effects of mPFC silencing, but inaccurately gives the impression that performance is at chance for food or water (PI ≈ 0, Extended Data Fig. 3), when the effect is strongly biased to food-seeking. For comparison, the raw data and behavioral summary of all animals (responders and non-responders) are reported in Extended Data Fig. 3 (see also Extended Data Table 1).

#### Analysis of trials with laser stimulation

For the pre-choice stimulation, we are comparing trials 25-50 and 75-100 of the stimulation session with the same trials (25-50 and 75-100) of the previous non-stimulation session in the same need state. For the reward evaluation period comparison, all 100 trials of the stimulation session were compared to all 100 trials of the prior non-stimulation session.

#### Analysis of conditioned place preference

The position of the mouse was tracked during all sessions. Based on the initial preference test, we calculated the percentage of time spent on each side and assigned each animal either the left or right side of the chamber, depending on which side was less preferred, where the animal would receive photostimulation. We then calculated the percentage of time spent on that side before any photostimulation (pre), after 5 consecutive days of passive conditioning (1^st^ post), during close-loop online stimulation (active) and the day after (2^nd^ post).

#### Analysis of mPFC silencing in animals switched from constant thirst to hunger

In Fig. 4b we compare mice (n=6) in thirst that were first switching between need states but then held constant in thirst for at least 5 training sessions over 10 consecutive days. The data of the same animals are compared without (regular) and with mPFC inhibition during the reward evaluation period during the end of the constant thirst training before those animals were switched to hunger. In Fig. 4c, we are comparing the performance of animals that were originally switching between need states, then held constant in thirst (for at least 5 training sessions and 10 consecutive days) and were then switched back to hunger. One group of animals (n=7) was tested in hunger without photostimulation (regular), while a different group of animals (n=7) was tested in hunger with photostimulation during the reward evaluation period.

#### Curve Fitting

To analyze and compare the intra-session learning curve of animals, we first excluded all trials lacking a choice from the data set and calculated the mean in a moving window of 3 trials to aid in fitting the Weibull function. We used a modified Weibull function^29^ to include an offset term (*a*) to fit the learning curves across trials (*t*) of each individual animal with the following function

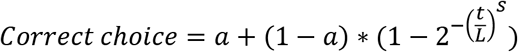

with parameters corresponding to offset (*a*), onset latency (*L*), and shape/steepness of function (*S*) fitted at trial t. We calculated the cumulative density function of those three fitted parameters and used the Kolmogorov-Smirnov test to detect difference between hunger and thirst. Curve fitting to the Weibull function was performed using the nlinfit function in Matlab.

#### Statistics

Data are reported as means ± s.e.m., unless otherwise stated. Pairwise comparisons were calculated by unpaired or paired nonparametric rank tests like Mann-Whitney U-test and Wilcoxon signed rank test, respectively, while learning curves were analyzed using ANOVA (see Extended Data Table 1 for details). All statistical tests were two-sided unless otherwise stated. For post hoc p-value correction we used Dunn’s test for multiple comparisons. Analyses were performed using Matlab (Mathworks).

---

**Extended Data Figure 1:**
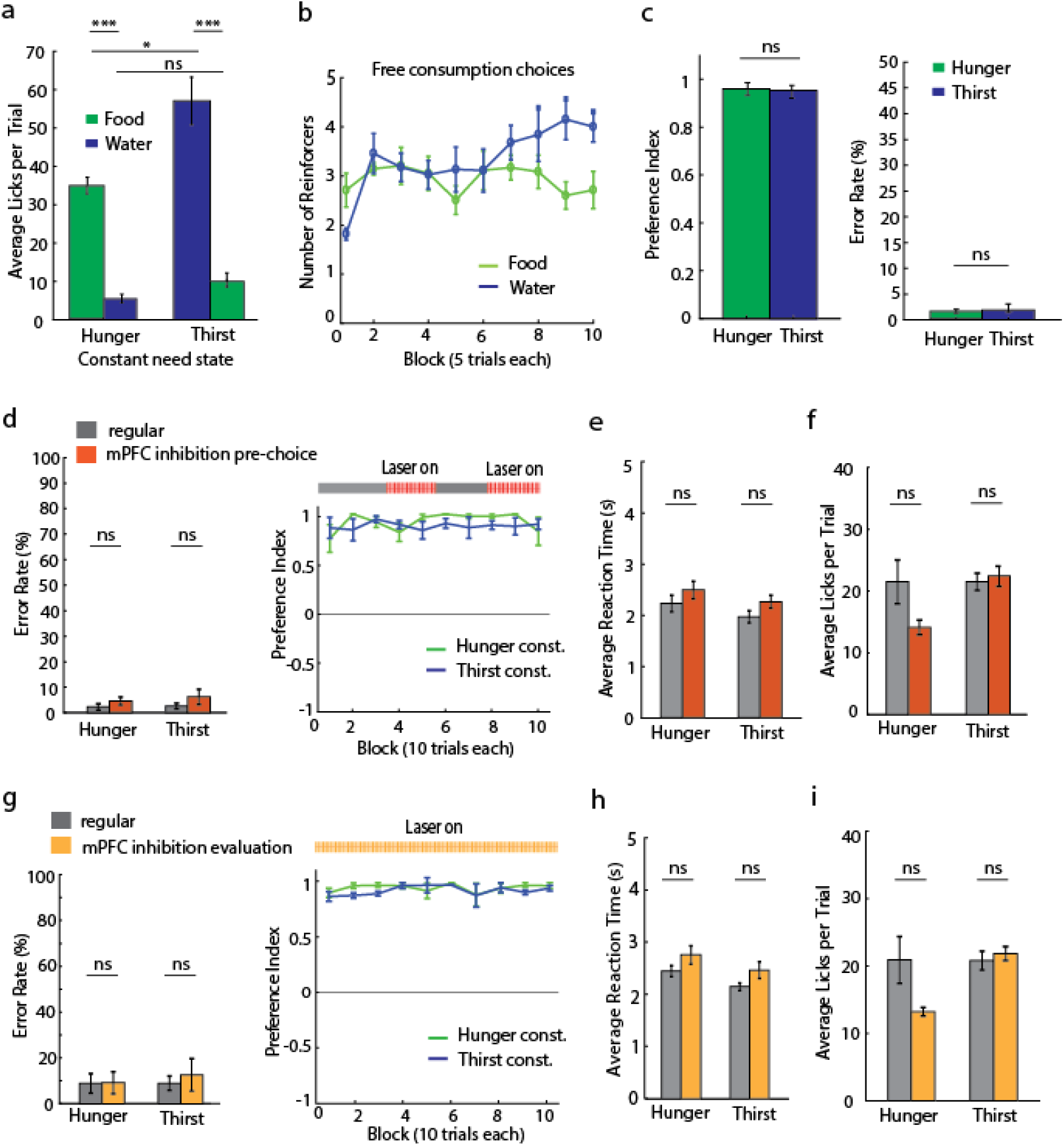
mPFC is not required for constant need-state decision-making. **a**, Lick-triggered food and water delivery preference test showing average licks per trial in hunger (n = 9) and thirst (n = 9) during constant need-state conditions. **b**, Comparison of gelled food and water reinforcer value in lick-triggered consumption test. Mice consumed a similar amount of water in thirst as they did for gelled food in hunger (n = 9 for each), indicating that the value of water in thirst and food in hunger was comparable under free consumption choice conditions. **c**, Preference index (left) and error rates (right) of lever press choices in constant need state (n = 9 in each hunger and thirst). **d-f**, Effect of mPFC silencing during pre-choice phase in constant need-state animals on error rate and preference index, reaction time of correct presses (**e**) and lick count (**f**) in hunger and thirst with (red) or without (grey) mPFC silencing. **g-i**, Effect of mPFC silencing during the reward evaluation period in constant need-state animals on error rate and preference index (**g**), reaction time of correct presses (**h**) and lick count (i) in hunger and thirst with (orange) or without (grey) mPFC silencing (regular n = 13, stimulation n =9 for each hunger and thirst). Error bars represent SEM. *P<0.05; **P<0.01; ***P<0.001; ns, P>0.05. Statistical analysis in Extended Data Table 1.

**Extended Data Figure 2:**
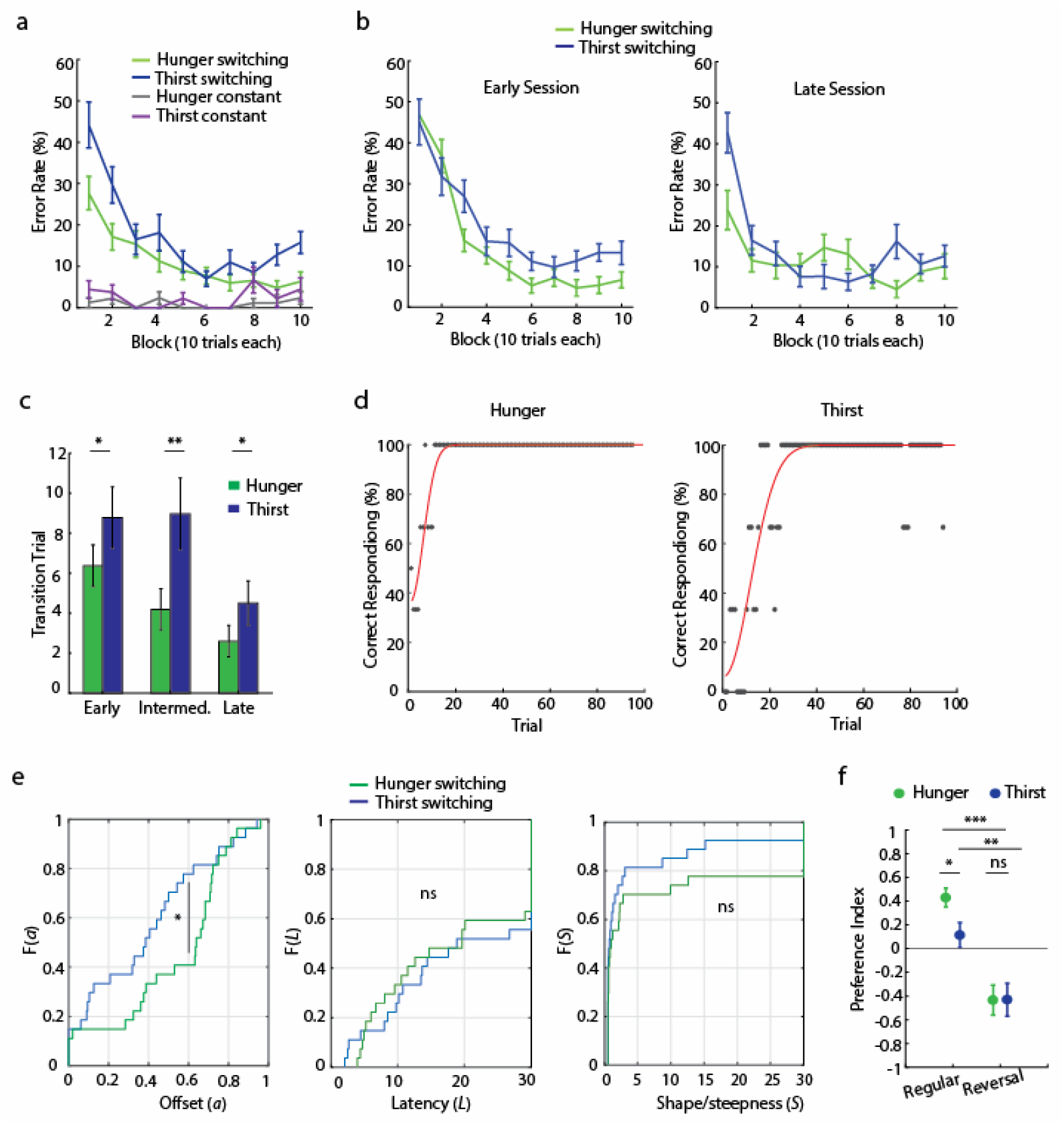
Food bias and learning rates for decision-making in thirst and hunger. **a**, Error rates for decisions in hunger and thirst with need-state switching (n = 27) for mice with intermediate experience on the task under need state-switching conditions as well as for mice during constant need state (n = 18). **b**, Error rates during early decision-making sessions under need state-switching conditions just after the task had been learned (left) and after extensive experience with task in late sessions (right) (early n = 27, late n = 22). **c**, Transition trial in hunger and thirst when correct performance exceeded 80% correct during early, intermediate and late sessions shows lag in correct performance in thirst due to initial food-seeking bias. **d**, Example of Weibull fit to correct responses for one session in hunger (left) and thirst (right). **e**, Cumulative density function for the three Weibull fit parameters (n = 27). (Left) offset *a* (*P<0.05), (middle) onset Latency *L*, (right) shape/steepness of function *S*. **f**, To test if mice determine which lever gives the greatest reward for their current need in each session, we reversed the lever contingencies (‘lever reversal’, n = 14). In both hunger and thirst, the initial performance was opposite to the reversed contingencies. Error bars represent SEM. *P<0.05; **P<0.01; ***P<0.001; ns, P>0.05. Statistical analysis in Extended Data Table 1.

**Extended Data Figure 3:**
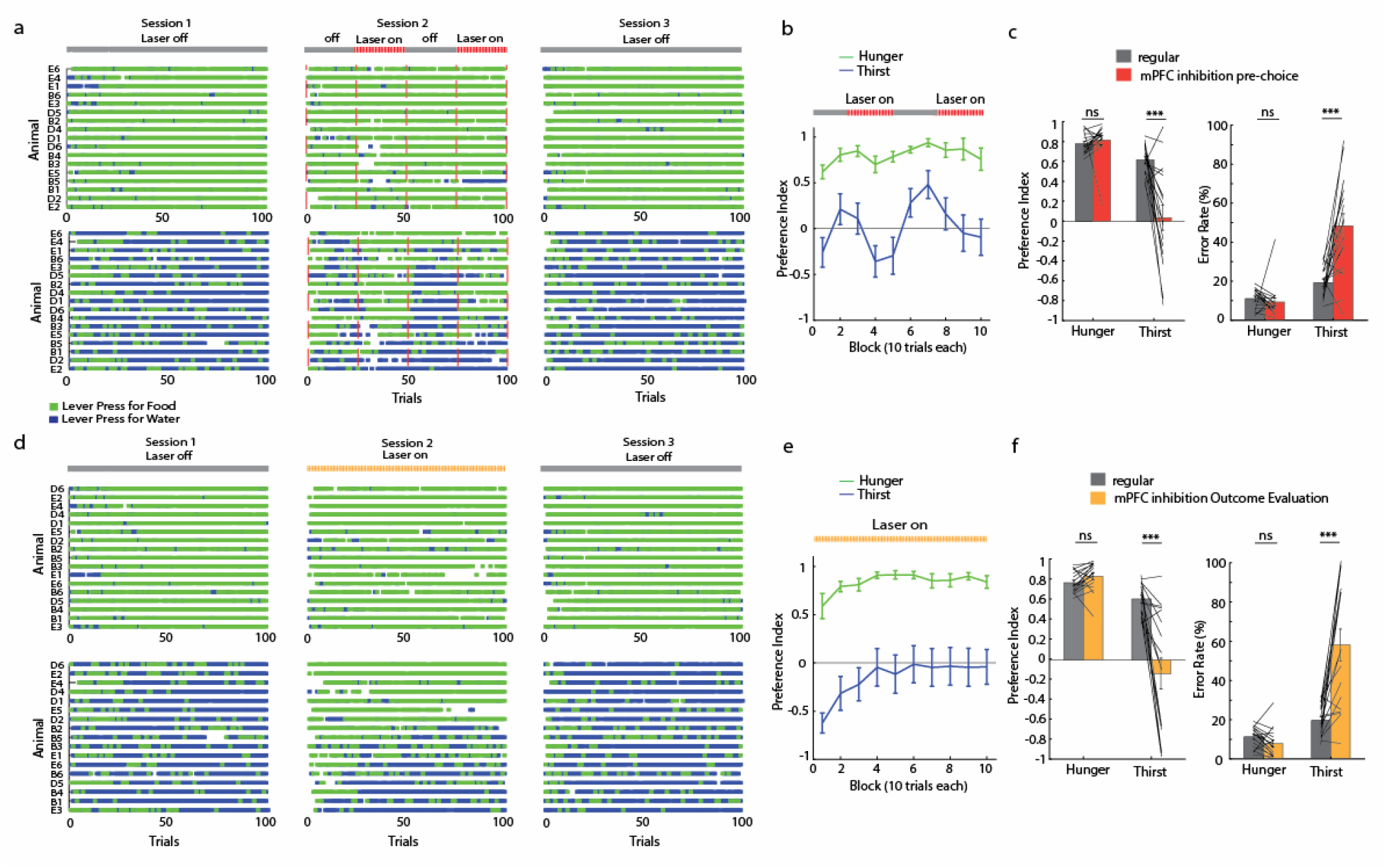
Population data for silencing medial prefrontal cortex. **a**, Individual animal performance (n = 17) before (session 1), during (session 2) and after (session 3) prechoice stimulation for hunger (upper panel) and thirst (lower panel). Tick marks are lever press for food (green) or water (blue). **b**, Preference index across trial blocks during pre-choice stimulation (n = 17 animals). **c**, Mean preference index and error rate during pre-choice mPFC silencing (n=17). **d**, Individual animal performance (n = 17) before (session 1), during (session 2) and after (session 3) reward evaluation stimulation for hunger (upper panel) and thirst (lower panel). **e**, Preference index across trial blocks during outcome evaluation stimulation (n=17). **f**, Mean preference index and error rate during outcome evaluation mPFC silencing. Error bars represent SEM. *P<0.05; **P<0.01; ***P<0.001; ns, P>0.05. Statistical analysis in Extended Data Table 1.

**Extended Data Figure 4.**
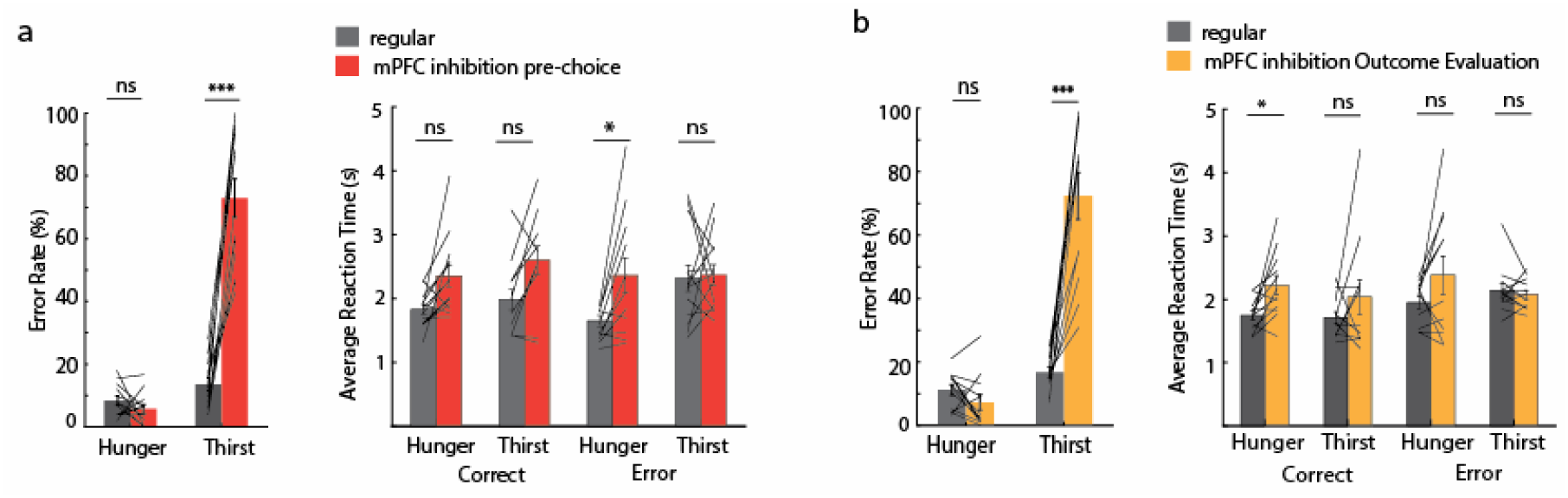
mPFC silencing during pre-choice and outcome evaluation. **a**, Error rates (left) and reaction times (right) for responders (see Methods) in the pre-choice stimulation period (n = 13). **b**, Error rates (left) and reaction times (right) for responders (see methods) in the reward evaluation period (n = 12). Error bars represent SEM. *P<0.05; **P<0.01; ***P<0.001; ns, P>0.05. Statistical analysis in Extended Data Table 1.

**Extended Data Figure 5:**
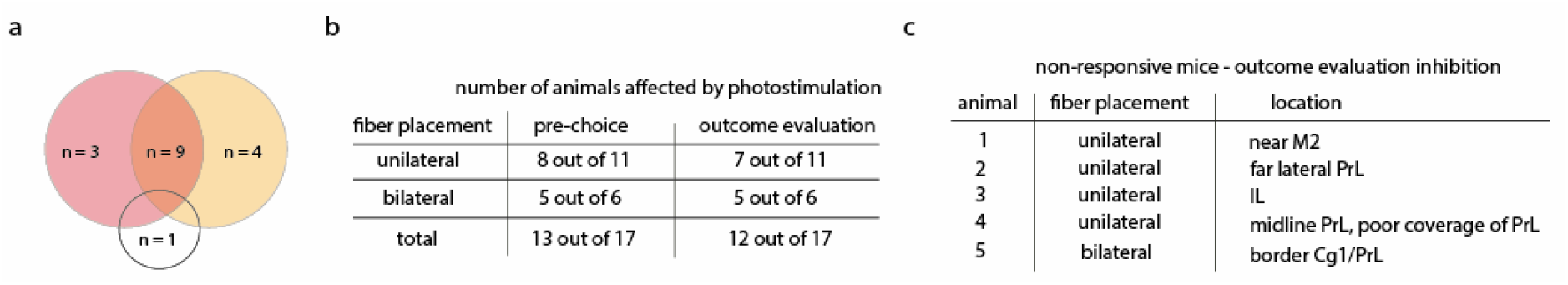
Details about responders vs. non-responders. **a**, Venn Diagram showing animals affected by mPFC silencing during the pre-choice (n = 13) and reward evaluation period (n = 12), with one animal being unaffected in both periods. **b**, Number of animals showing an effect in pre-choice and reward evaluation stimulation depending on uni-vs. bilateral fiber implant. **c**, Anatomical location of non-responding animals during outcome evaluation stimulation.

**Movie 1**. Example session of a thirsty animal showing lever press performance in trials before, during, and after optogenetic pre-choice mPFC silencing. After a variable inter-trial interval, each trial begins with the presentation of an auditory cue (1 s) which is presented followed by levers extension. In mPFC silencing trials, laser stimulation starts with the onset of cue presentation and lasts until the animal presses a lever or until the response window ends (5 s).

**Extended Data Table 1.** Results of statistical analyses.

| Figure | Sample size (n, mice) | Statistical Test | Values |
| --- | --- | --- | --- |
| 1d | Hunger: 9 \| Thirst: 9 | Mann-Whitney U Test:<br>*Photostim vs Regular* | Hunger Food: $P=0.29$ \| Water: $P=0.24$<br>Thirst Food: $P=1$ \| Water: $P=0.87$ |
| 1f | Constant Hunger:9<br>Thirst:9<br>Switch.<br>Hunger/Thirst:27 | Mann-Whitney U Test | Constant: $P=0.4$<br>Switching: $P=0.04$ |
| 1g | Hunger reg: 13 stim: 9<br>Thirst reg: 13 stim: 9 | Wilcoxon Signed Rank Test:<br>*Photostim vs Regular* | Pre-choice Hunger: $P=0.35$<br>Thirst $P=0.77$<br>Evaluation Hunger: $P=0.74$<br>Thirst: $P=0.94$ |
| 2c | 27 | 2-way rm ANOVA;<br>Factor 1: *Hunger vs Thirst*<br>Factor 2: *Block*<br>Interaction: *State x Block*<br>Post hoc p-value correction with<br>Dunn's test for multiple<br>comparisons | $F(1,234)=10.39, P=0.006$<br>$F(9,234)=23.91, P<0.001$<br>$F(9,234)=1.6, P=0.11$ |
| 2d | 27 | Wilcoxon signed rank test | Preference index: $P=0.01$<br>Error rate: $P=0.01$ |
| 2e | 27 | 2-way rm ANOVA;<br>Factor 1: *Hunger vs Thirst*<br>Factor 2: *Block*<br>Interaction: *State x Block* | $F(1,234)=14.1, P<0.001$<br>$F(9,234)=9.3, P<0.001$<br>$F(9,234)=1.85, P=0.059$ |
| 2f | 27 | Wilcoxon signed rank test:<br>*Correct Hunger vs Thirst*<br>*Error Hunger vs Thirst*<br>*Correct overall vs Error overall*<br>Post hoc p-value correction with<br>Dunn's test for multiple<br>comparisons | $P=0.004$<br>$P=0.1$<br>$P=0.0003$ |
| 2g | 27 | Wilcoxon Signed Rank Test:<br>*Hunger vs Thirst* | $P=0.015$ |
| 2h | Early: 27\| Interm:27 \|<br>Late:22 | Wilcoxon signed rank test<br>Post hoc p-value correction with<br>Dunn's test for multiple<br>comparisons | Early: $P=0.09$ \| Intermediate: $P=0.009$<br>Late: $P=0.014$ |
| 2i | Early: 27\| Interm:27 \|<br>Late:22<br>Constant: 18 | Wilcoxon signed rank test:<br>*Hunger vs Thirst*<br>Const: Mann-Whitney U Test:<br>*Hunger vs Thirst*<br>Post hoc p-value correction with<br>Dunn's test for multiple<br>comparisons | Early: $P=0.83$ \| Intermediate: $P=0.03$<br>Late: $P=0.008$<br>Const.: $P=0.73$ |
| 2j | Regular switching:27<br>After Constant Thirst:<br>7 | Mann-Whitney U Test<br>*Block 1, Block 2*<br>Post hoc p-value correction with<br>Dunn's test for multiple<br>comparisons | $P=0.042$<br>$P=0.031$ |
| 2k | Hunger: 6 \| Thirst: 7 | 2-way rm ANOVA separate for Hunger & Thirst<br>Factor 1: *before vs pre-exposure*<br>Factor 2: *Block*<br>Interaction (IA): *Condition x Block*<br>Post hoc multiple comparison: Tukey's HSD test.<br>*Thirst Block 1, Hunger Block 1* | Thirst Factor 1: $F(1,54)=11.16, P=0.015$<br>Thirst Factor 2: $F(9,54)=3.5, P=0.0017$<br>Thirst IA: $F(9,54)=1.98, P=0.058$<br>Hunger Factor 1: $F(1,45)=1.8, P=0.23$<br>Hunger Factor 2: $F(9,45)=12.91, P<0.001$<br>Hunger IA: $F(9,45)=2.96, P=0.007$<br>$P<0.01, P<0.05$ |
| 3c | 13 | Wilcoxon signed rank test:<br>*Photostim vs Regular*<br>Post hoc p-value correction with Dunn's test for multiple comparisons | Hunger: $P=0.24$<br>Thirst: $P<0.001$ |
| 3d | 13 | Wilcoxon signed rank test:<br>*Photostim vs Regular* | Hunger correct: $P=0.78$ \| error: $P=0.53$<br>Thirst correct: $P=0.76$ \| error: $P=0.89$ |
| 3g | 12 | Wilcoxon signed rank test:<br>*Photostim vs Regular*<br>Post hoc p-value correction with Dunn's test for multiple comparisons | Hunger: $P=0.15$<br>Thirst: $P<0.001$ |
| 3h | 12 | Wilcoxon signed rank test:<br>*Photostim vs Regular*<br>Post hoc p-value correction with Dunn's test for multiple comparisons | Hunger correct: $P=0.56$ \| error: $P=0.76$<br>Thirst correct: $P=0.01$ \| error: $P=0.06$ |
| 3i | 4 | Wilcoxon signed rank test:<br>*Pre vs Post1; Pre vs Active; Active vs Post2*<br>Post hoc p-value correction with Dunn's test for multiple comparisons | $P=0.99$<br>$P=0.56$<br>$P=0.96$ |
| 4b | 6 | Wilcoxon signed rank test:<br>*Photostim vs Regular* | Preference Index $P=0.15$<br>Error Rate $P=0.15$<br>RT correct $P=0.16$ \| error $P=0.63$<br>Licks correct $P=0.84$ \| error $P=0.69$ |
| 4c | 7 | Mann-Whitney U Test:<br>*Photostim vs Regular* | Preference Index $P=0.016$<br>Error Rate $P=0.016$<br>RT correct $P=0.01$ \| error $P=0.07$<br>Licks correct $P=0.77$ \| error $P=0.36$ |
| ED1a | 9 | Mann-Whitney U Test<br>Post hoc p-value correction with Dunn's test for multiple comparisons | Hunger/Food vs Hunger/Water: $P<0.001$<br>Thirst/Food vs Thirst/Water: $P=0.0009$<br>Hunger/Food vs Thirst/Water: $P=0.02$<br>Hunger/Water vs Thirst/Food: $P=0.08$ |
| ED1b | 9 | 2-way rm ANOVA; Factor 1:<br>*Food vs Water*<br>Factor 2: *Block*<br>Interaction: *Reward x Block* | $F(1,72)=5.06, P=0.28$<br>$F(9,72)=2.62, P=0.01$<br>$F(9,72)=1.8, P=0.07$ |
| ED1c | Hunger: 9 \| Thirst: 9 | Wilcoxon signed rank test | Preference Index: $P=0.15$<br>Error Rate: $P=0.15$ |
| ED1d | Reg: 13 \| stim: 9 | Wilcoxon signed rank test:<br>*Photostim vs Regular* | Hunger: $P=0.35$ \| Thirst: $P=0.77$ |
| ED1e | Reg: 13 \| stim: 9 | Wilcoxon signed rank test:<br>*Photostim vs Regular* | Hunger: $P=0.31$ \| Thirst: $P=0.12$ |
| ED1f | Reg: 13 \| stim: 9 | Wilcoxon signed rank test:<br>*Photostim vs Regular* | Hunger: $P=0.12$ \| Thirst: $P=0.54$ |
| ED1g | Reg: 13 \| stim: 9 | Wilcoxon signed rank test:<br>*Photostim vs Regular* | Hunger: $P=0.68$ \| Thirst: $P=1$ |
| ED1h | Reg: 13 \| stim: 9 | Wilcoxon signed rank test:<br>*Photostim vs Regular* | Hunger: $P=0.12$ \| Thirst: $P=0.12$ |
| ED1i | Reg: 13 \| stim: 9 | Wilcoxon signed rank test:<br>*Photostim vs Regular* | Hunger: $P=0.1$ \| Thirst: $P=0.35$ |
| ED2c | Early: 27 \| Interm:27 \|<br>Late:22 | Wilcoxon signed rank test<br>Post hoc p-value correction with<br>Dunn's test for multiple<br>comparisons | Early: $P=0.045$<br>Intermediate: $P=0.009$<br>Late: $P=0.015$ |
| ED2f | 27 | Kolmogorov-Smirnov test | $\alpha$: $P<0.05$ \| $L$: $P=0.99$ \| $S$: $P=0.69$ |
| ED2g | Regular: 27 \|<br>Reversal: 14 | Mann-Whitney U Test:<br>*Regular Hunger vs Thirst;*<br>*Reversal Hunger vs Thirst;*<br>*Regular Hunger vs. Reversal*<br>*Hunger;*<br>*Regular Thirst vs Reversal Thirst*<br>Post hoc p-value correction with<br>Dunn's test for multiple<br>comparisons | $P=0.04$<br>$P=0.98$<br>$P<0.0001$<br>$P=0.0057$ |
| ED3c | 17 | Wilcoxon signed rank test:<br>*Photostim vs Regular* | Preference Index: Hunger: $P=0.07$<br>Thirst: $P=0.001$<br>Errors: Hunger: $P=0.07$<br>Thirst: $P=0.001$ |
| ED3f | 17 | Wilcoxon signed rank test:<br>*Photostim vs Regular* | Preference Index: Hunger: $P=0.08$<br>Thirst: $P=0.0007$<br>Errors: Hunger: $P=0.08$<br>Thirst: $P=0.0007$ |
| ED4a | 13 | Wilcoxon signed rank test:<br>*Photostim vs Regular*<br>Post hoc p-value correction for<br>reaction times with Dunn's test<br>for multiple comparisons | Errors, Hunger: $P=0.24$<br>Errors, Thirst: $P<0.0002$<br>Reaction Time, Hunger correct:<br>$P=0.096$<br>Hunger, error: $P=0.032$<br>Reaction Time, Thirst correct: $P=0.054$<br>Thirst, error: $P=0.78$ |
| ED4b | 12 | Wilcoxon signed rank test:<br>*Photostim vs Regular*<br>Post hoc p-value correction for<br>reaction times with Dunn's test<br>for multiple comparisons | Errors, Hunger: $P=0.15$<br>Errors, Thirst: $P=0.0004$<br>Reaction Time, Hunger correct:<br>$P=0.024$<br>Hunger, error: $P=0.46$<br>Reaction Time, Thirst correct: $P=0.17$<br>Thirst, error: $P=0.56$ |

